# The mechanical properties of *Arabidopsis thaliana* roots adapt dynamically during development and to stress

**DOI:** 10.1101/2025.07.29.667350

**Authors:** Luis Alonso Baez, Astrid Bjørkøy, Francesco Saffioti, Sara Morghen, Dhika Amanda, Michaela Tichá, Maarten Besten, Anastasiia Ivanova, Joris Sprakel, Bjørn Torger Stokke, Thorsten Hamann

## Abstract

Mechanical properties of plant cells and tissues change dynamically, influencing plant growth, development, and interactions with the environment. Despite their central roles in plant life, current knowledge of how these properties change *in vivo* is very limited. Here we have combined Brillouin microscopy and molecular rotors to investigate stiffness, viscosity and porosity in living *Arabidopsis thaliana* seedling roots during differentiation and in response to stress and genetic manipulation. We found that mechanical properties change in a cell- and tissue-specific manner. The properties change dynamically during differentiation to support directional cell expansion. Cell-type-specific adaptations are induced within hours in response to stress or changes in cell wall metabolism. The findings form the foundation for future studies to characterize regulatory mechanisms linking biochemical signaling and mechanical properties.

## Main Text

The mechanical properties of plant structures, ranging from the sub-cellular to the organ level, are key determinants for plant growth, development, and interactions with the environment (*1–4*). These properties arise from the interplay between turgor pressure, molecular crowding, and sub-cellular structures such as the cytoskeleton, cell wall, or nuclear envelope (*5–7*). Their dynamic nature is tightly linked to biological function, yet our understanding of how mechanical properties vary across tissues and scales remains limited (*8–10*). In particular, sub-epidermal plant cell layers have been characterized predominantly through indirect methods while *in vivo* measurements have largely been restricted to surface tissues or whole organ analyses (*11–15*).

Recently, non-invasive technologies have become available enabling *in vivo* analyses of subepidermal mechanical properties with subcellular and temporal resolution. Two complementary approaches exemplify these advances: Brillouin microscopy and BODIPY-based molecular rotors (*16–18*). Molecular rotors report on local mechanical properties through environmentally driven changes in rotor configuration, which affect its fluorescence lifetime. In this study, we used CarboTag dyes to assess hydrodynamic porosity, reflecting the space available for probe rotation within the cell wall matrix (*19*). Brillouin microscopy provides label-free measurements of stiffness and viscosity by detecting frequency shifts and changes in the linewidth of the inelastically scattered light from the sample, with increases in frequency shift or linewidth implying increased stiffness and viscosity (*20*). Stiffness is related to material deformation or compression, while viscosity is a resistance to flow or movement between adjacent layers. Frequency shifts can be used to obtain the longitudinal storage modulus, describing the ratio of uniaxial strain to uniaxial stress. Linewidths are related to the loss modulus, reflecting viscous damping and energy dissipation within the material. The calculation of these moduli requires that the refractive index and mass density of the material are known. Brillouin microscopy, unlike Young’s modulus measurements which produce values in the MPa range, captures mechanical behavior at GHz frequencies, yielding higher moduli in the GPa range. Importantly, stiffness and viscosity changes are not necessarily correlated, highlighting the value of measuring both parameters to dissect complex mechanical properties (*21*, *22*).

Simultaneous analysis of multiple mechanical properties is essential to fully understand the molecular processes controlling them in plant tissues and cell types during development and environmental interactions. This approach opens new avenues to elucidate causal relationships between mechanical patterns observed at the subcellular level with those detected at tissue and organ levels (*8*, *10*, *23*). Here we begin to provide these insights by generating *in vivo* maps of the mechanical properties of *Arabidopsis thaliana* (Arabidopsis) root layers along developmental axes and examining how the properties change during differentiation with single cell resolution, in response to stress and to genetic alterations in cell wall metabolism and signaling. These results establish a framework to investigate the root mechanical properties across spatial scales, developmental stages and environmental interactions, laying the groundwork for uncovering regulatory mechanisms and developing mechanobiological *in vivo* models (*24–27*).

### Tissue-specific mechanical maps of Arabidopsis root tips

Tissues in Arabidopsis seedling roots are organized as radially concentric cylindrical layers (epidermis, cortex, endodermis, and stele). Because of their different functions in development and stress responses, it has been postulated that individual tissue layers have distinct mechanical properties (*28*, *29*). However, this has never been experimentally validated *in vivo* in a non-invasive manner because direct *in vivo* measurements of mechanical properties in subepidermal tissues have been technically challenging and mostly relied on indirect observations (*4*, *10*, *11*, *13*).

To close this major knowledge gap, we characterized the four main tissue types in 6-day-old Arabidopsis seedling roots, using confocal and Brillouin microscopy. First, we obtained fluorescence images of Col-0 roots expressing the p35S:LTI6b-GFP plasma membrane marker (Fig. 1A). Subsequently, we scanned the same areas to establish the local mechanical properties (Fig. 1B and 1C). For each tissue, we quantified the spatial profile of the frequency shift (related to stiffness) and the linewidth (related to viscosity) along the developmental axis of the root, from the meristem zone above the stem cell niche to the end of the elongation zone (Fig. 1D).

**Fig. 1.**
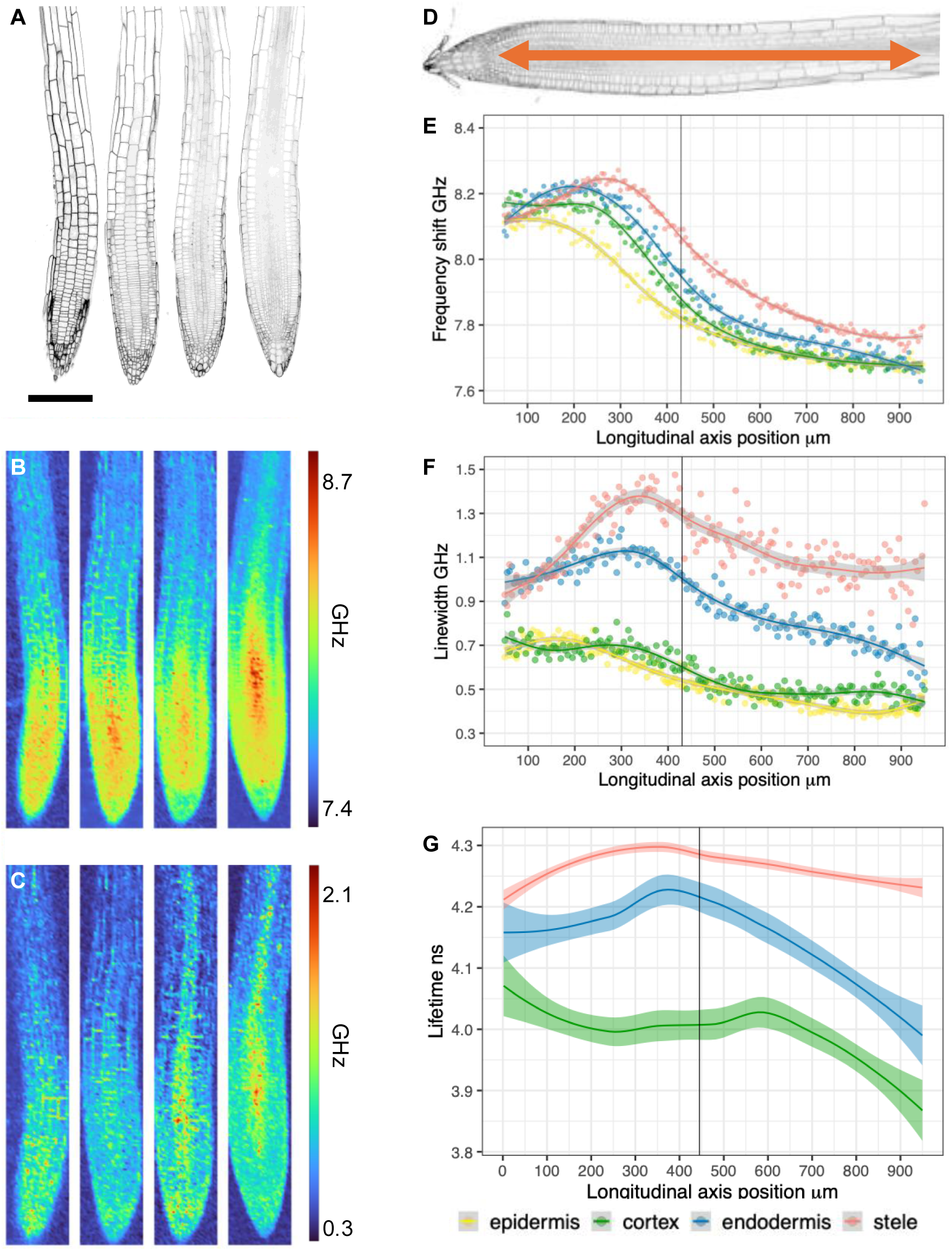
Maps of the mechanical properties of the Arabidopsis root tip. (A and B) Left to right, image of epidermis, cortex, endodermis and stele planes. (A) Confocal images using p35S:LTI6b-GFP plasma membrane marker. Scale bar 150 µm. (B) Frequency shift and (C) linewidth heatmaps, 150 x 1000 µm, of the areas scanned in A. (D) Direction of quantification along the longitudinal root axis. (E) Frequency shift, (F) linewidth and (G) cell wall porosity quantification.The vertical line indicates the average position of the transition zone identified with respect to the cortex file.

The stele plane showed a unique frequency shift profile (Fig. 1E). While its values in the vicinity of the stem cell niche were comparable to those of the other tissues, the frequency shift increased markedly in the central meristem. It remained elevated relative to the surrounding tissues before decreasing just prior to the transition zone and dropping further in the elongation zone. Frequency shift levels in the epidermis, cortex, and endodermis layers exhibited similar values in the meristematic region. The position of the maximum frequency shifts differed between the tissues. The maximum frequency shift was located near the stem cell niche in the epidermis, in the middle region of the meristem for both cortex and endodermis and a few cells before the transition zone in the stele (defined as the position in the cortex file where cells start to elongate) (*30*). The epidermis values gradually declined towards the elongation zone, whereas cortex and endodermis values increased slightly in the meristem before declining. Among the three tissues, the epidermis consistently exhibited the lowest frequency shift values, while cortex and endodermis remained higher. Linewidth measurements in the stele were markedly higher than in the other three tissues along the developmental axis (Fig. 1F). The lowest values were observed near the stem cell niche, followed by a gradual increase that peaked in the central meristem before decreasing toward the elongation zone. In the endodermis, linewidth values began at levels comparable to the stele near the stem cell niche but exhibited only a modest increase in the meristem, followed by a decrease toward the transition and elongation zones. The cortex and epidermis showed similar patterns, with consistently lower linewidths than the stele and endodermis. Although the changes were more subtle, both displayed a gradual decline from the meristem to the elongation zone.

To validate these findings, we visualized local changes in cell wall porosity using CarboTag-BDP molecular rotors (*19*). Median plane profiles of roots revealed elevated fluorescence lifetimes in the meristem zone relative to the elongation zone (Supp. Fig. 1A). We quantified the longitudinal porosity profile of cell walls along the developmental axis for distinct root tissues (the epidermal layer was not quantified due to chemical quenching of the probe at the root surface (*19*)) (Fig. 1G). In both stele and endodermis layers, lifetime values increased from the stem cell niche to the end of the meristem zone and then declined to lower values in the elongation zone. The lifetime values in the cortex layer were higher in the stem cell niche, decreased to a plateau in the meristem zone and decreased further in the elongation zone. These results are consistent with our Brillouin-based measurements and confirm that mechanical properties change dynamically along the longitudinal root axis.

Our measurements identified distinct and tissue-specific profiles associated with stiffness and viscosity in the seedling root *in vivo*. The patterns suggest that cells undergo dynamic changes in their mechanical properties during the developmental transition from division to elongation, a process that has been reported to take between one (*31*, *32*) to three days (*33*). Stiffness, viscosity and porosity exhibit similarities along the developmental axes but also show differences, highlighting the importance of investigating more than one mechanical property. These observations imply the existence of regulatory mechanisms modulating mechanical properties with single-cell resolution.

### Mechanical properties of single cells

As differentiation progresses, anisotropic mechanical properties between transversal and longitudinal cell walls are thought to be modified in a tightly controlled manner to allow cell expansion along the longitudinal axis (*14*, *18*, *34*). We chose the cortex layer as a model to investigate *in vivo* the temporal and spatial dynamics of mechanical properties in individual cells along the developmental axis (Fig. 2A). We established and compared mechanical properties of longitudinal and transversal cell walls in the cortex cell layer in three different developmental stages (Fig. 2B-E). In the meristematic zone, frequency shifts were similar in both longitudinal and transversal walls, while linewidth was higher in the transversal walls. In the transition zone, both frequency shift and linewidth values were higher in transversal walls compared to longitudinal ones. In the elongation zone, frequency shifts in transversal cell walls were increased compared to longitudinal cell walls while no significant differences in the linewidth existed between them. The frequency shift values of the cortex cell walls decreased from the meristem to the elongation zone, in agreement with the results of the global map of the root tip. Notably, the decrease was more pronounced in the longitudinal walls compared to the transversal walls, which is consistent with the direction of cell elongation along the developmental axis. We also detected heterogeneous mechanical properties in individual cell walls with non-uniform frequency shift and linewidth values, suggesting local variation in mechanical properties possibly reflecting locations where new cell wall materials are being synthesized and/or deposited (Figures 2B and 2D, transition and elongation zones). In complementary experiments using CarboTag-BDP, transverse cell walls exhibited higher porosity than longitudinal walls across all developmental stages examined (Supp. Fig. 1B). To summarize, these results suggest that mechanical anisotropy may already be present in the meristem zone because viscosity and porosity seemed notably higher in transverse cell walls while stiffness differences were not significant. This would be consistent with expansion being limited while cell division activity is ongoing (*35*). In the transition and elongation zones, longitudinal cell walls were less stiff than transversal cell walls, allowing directional elongation as predicted by theoretical models and previous experiments (*14*, *23*, *34*).

**Fig. 2.**
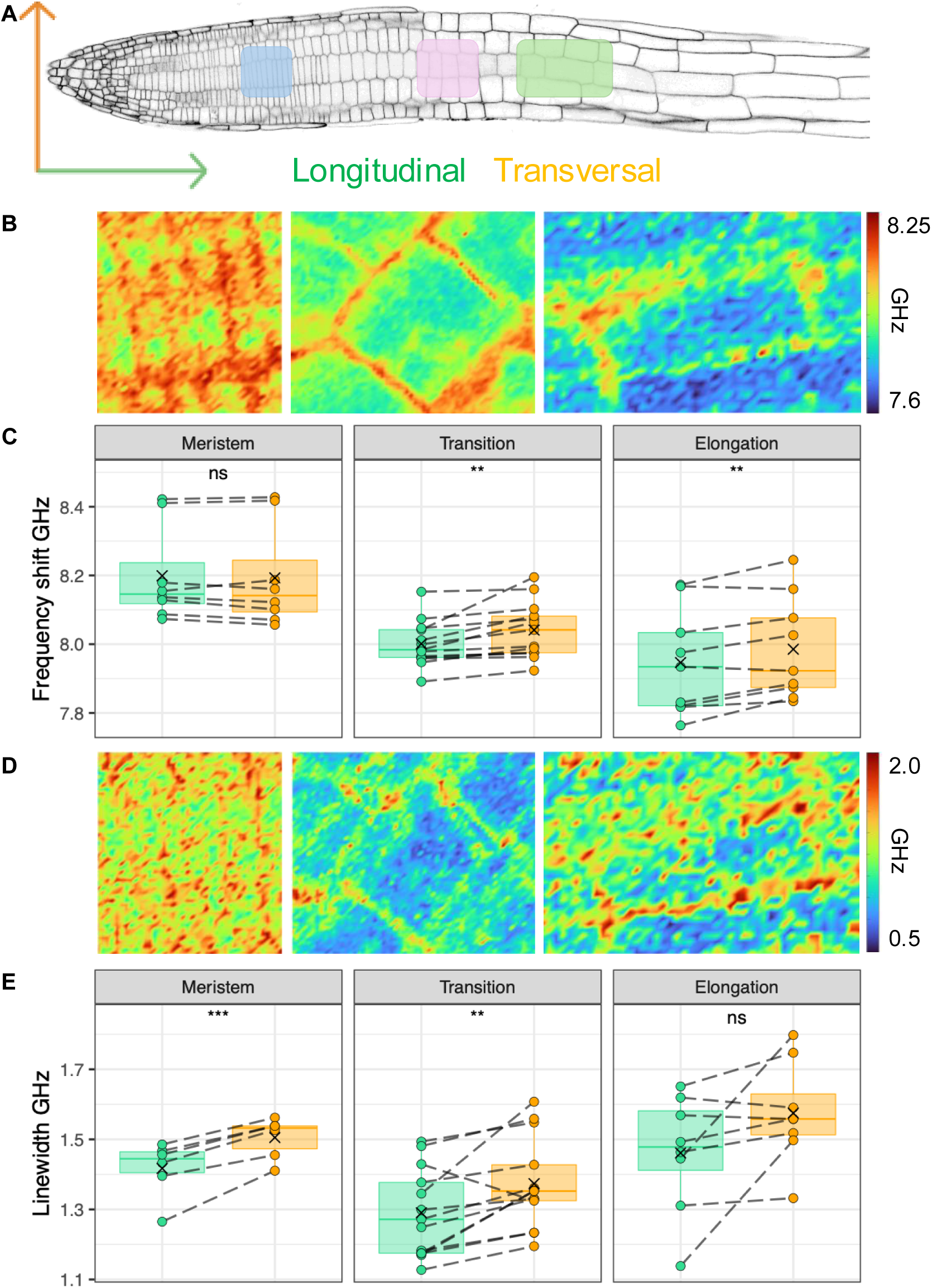
Anisotropic mechanical properties of Arabidopsis cell walls in the cortex layer. (A) Orientations of cell walls and confocal image of the cortex plane vizualized with p35S:LTI6b-GFP. Regions where imaging was performed are indicated for the meristem (blue), transition zone (pink) and elongation zone (green). (B) Heatmaps of the frequency shift in cells at the meristem, 50 x 40 µm (left), transition zone, 55 x 60 µm (middle) and elongation zone, 35 x 50 µm (right). (C) Quantification of the frequency shifts at cell walls. (D) Heatmaps of the linewidth in cells at the meristem (left), transition zone (middle) and elongation zone (right). (E) Quantification linewidth at cell walls.

To investigate spatial variation in stiffness and viscosity within an individual cell further, we selected root hair cells, since they form a well-characterized and representative model for polarized cell expansion (*36*, *37*) (Fig. S2A). The p35S:LTI6b-GFP reporter was used as an indicator for plasma membrane integrity and root hairs, ranging in length from 30 to 65 μm, were analyzed (Fig. S2B-C). In 5 out of 19 cells we detected an ellipsoidal region with elevated frequency shift and linewidth, probably representing the nucleus (Fig. S2D). We quantified the frequency shift and linewidth in three regions – tip, middle and base, excluding the nuclear region when present (Fig. S2A). The frequency shift was significantly higher at the base compared to the middle and tip, whereas the linewidth did not differ between the three regions, suggesting that viscosity is conserved while stiffness differs between the observed areas (Fig.S2E). In parallel, we measured the refractive index (*n*) of the root hair cell peripheries using quantitative phase imaging (Supp. Fig. 2F), obtaining values consistent with previous reports (average *n* = 1.38) (*38*). Based on these measurements, we calculated the corresponding mass density (see Methods) (*39*) and, found that the longitudinal modulus at the root hair periphery ranged from 2.62 to 3.07 GPa, values typical of soft biological materials (*40*). These values exceed the MPa-range reported by atomic force microscopy, due to higher frequency operation of Brillouin microscopy (*41*). These results provide insights into the *in vivo* mechanical properties of root hairs, quantifying the longitudinal modulus and supporting observations made above for cortex cells, namely that subcellular differences in mechanical properties exist.

### Mechanical properties differ between tissue layers in the elongation zone

Root cells stop dividing and start their directional growth in the elongation zone, marking a fundamental switch in differentiation. This developmental transition is particularly sensitive to perturbations, leading to alterations in the morphology of the cell wall or plasma membrane occurring in this region first. To investigate the impact of perturbations on mechanical properties in individual cell types, we first characterized the mechanical properties of cell walls and cytoplasm in the various tissue layers in the middle part of the elongation zone under control conditions (Fig. 3A-C). We found that frequency shifts in the outer epidermal cell walls were lower than those of all other tissues (Fig. 3D). Interestingly, the cell wall separating epidermis from cortex showed the highest mean frequency shift value of all cell walls examined. The remaining inner cell walls also exhibited shifts higher than the outer epidermal cell wall. The linewidth changed in a stepwise manner, with the lowest values exhibited also by the outer-epidermal wall, intermediate values in the epidermis-cortex, cortex-endodermis and endodermis-pericycle walls and the highest values detected in the stele. In both epidermis and cortex cells, we regularly observed cytoplasmic areas with increased linewidth values compared to the remainder of the respective cell. The structures are at least 10 μm in length (Fig. 3C). In the cytoplasm, frequency shift and linewidth changed in a similar manner (Fig. 3E). Lowest values were observed in the epidermis with a gradual increase towards inner tissues and a peak in the stele. These observations suggest pronounced differences in the mechanical properties of outer epidermal versus inner cell walls and a more gradual change in the properties of the cytoplasm towards the stele.

**Fig. 3.**
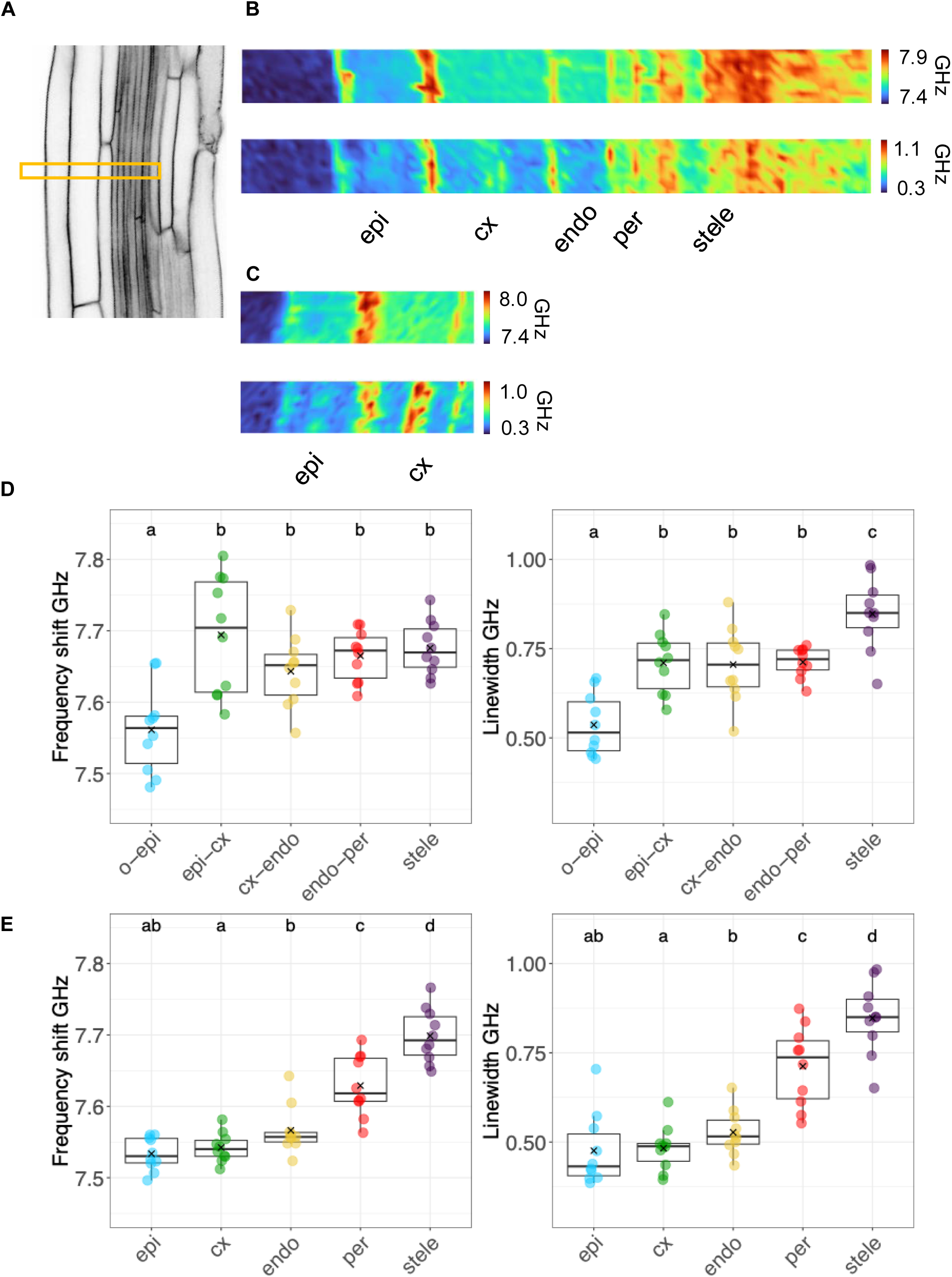
Mechanical properties of cell walls and cytoplasm in the elongation zone. (A) Confocal image of the root elongation zone median plane with p35S:LTI6b-GFP. Yellow rectangle indicates the area scanned with the Brillouin microscope. (B) Heatmaps (10×100 µm) of frequency shift (top) and linewidth (bottom). (C) Focused heatmaps (10×42 µm) of epidermis and cortex cells showing frequency shift (upper) and linewidth (lower). (D) Quantification of the frequency shift (left) and linewidth (right) of cell walls. (E) Quantification of the frequency shift (left) and linewidth (right) of the cytoplasm. epi= epidermis, cx = cortex, endo = endodermis, per = pericycle.

We also examined 4-day-old seedlings to determine if mechanical properties change with developmental age. Frequency shifts and linewidths in both cell walls and cytoplasm displayed qualitatively similar trends as observed in older seedlings. Outer tissues layers exhibited the lowest and the stele the highest values (Supp. Fig, S3A-D), suggesting that mechanical properties of the central elongation zone are stable in this time-window.

### Perturbations in cell wall biosynthesis and integrity affect mechanical properties

We examined next if genetic manipulation of cell wall metabolism and signaling affect the mechanical properties of the elongation zone in a specific manner using the following mutants: *procuste1* (*prc1-1*), containing a point mutation in *CELLULOSE SYNTHASE3* that causes reductions in cellulose synthesis (70% less cellulose) in roots (*42*) and dark-grown hypocotyls (*43*); *isoxaben resistant1* (*ixr1-1*), containing a point mutation in *CELLULOSE SYNTHASE6* leading to 30% less crystalline cellulose (*44*); *pectin methylesterase3* (*pme3*), impaired in demethylesterification of homogalacturonan and affecting mechanical strength of cell walls (*45*); and the double mutant *xylosyltransferase1, 2* (*xxt1xxt2*), defective in xyloglucan synthesis (*46*). In addition, we investigated *theseus1* (*the1-1*) and *feronia* (*fer-4*) seedlings, which have defects in cell wall integrity signaling and responses to mechanical stimuli (*47–50*). We also included *abscisic acid deficient2* (*aba2-1*) seedlings, which produce less abscisic acid (ABA), a hormone required to regulate turgor pressure in response to hyperosmotic stress (*50*). Brillouin measurements revealed tissue-specific alterations in mechanical properties of the cell walls and cytoplasm in certain mutants (Fig. 4 and Supp. Fig. 4A-C). Compared to the corresponding cell walls in Col-0, *xxt1xxt2* and *the1-1* exhibited lower frequency shifts in epidermis-cortex walls; *xxt1xxt2, the1-1,* and *pme3* in cortex-endodermis walls; *the1-1* in endodermis-pericycle walls while *pme3*, *xxt1xxt2*, and *the1-1* exhibited changes in the stele walls (Fig. 4A). *fer-4* exhibited an elevated frequency shift only in the stele walls, suggesting limited impact on mechanical properties. Linewidth values were reduced in endodermis-pericycle walls of *prc1-1,* and in the stele walls for *prc1-1* and *fer-4* (Fig. 4B). In the cytoplasm, reduced frequency shift values were detected in the epidermis in *pme3*, *xxt1xxt2* and *the1-1*; in the cortex in *pme3*, *xxt1xxt2*, *the1-1* and *aba2-1*; in the endodermis and pericycle in *the1-1*; and in the stele in *pme3*, *xxt1xxt2* and *the1-1*, compared to Col-0 controls (Fig.4C). Similar to corresponding cell walls, *fer-4* seedlings showed increased cytoplasmic frequency shifts in the stele. Cytoplasmic linewidths were reduced in the cortex and pericycle in *prc1-1* and in *prc1-1* and *fer-4* in the stele, compared to Col-0 (Fig.4D).

**Fig. 4.**
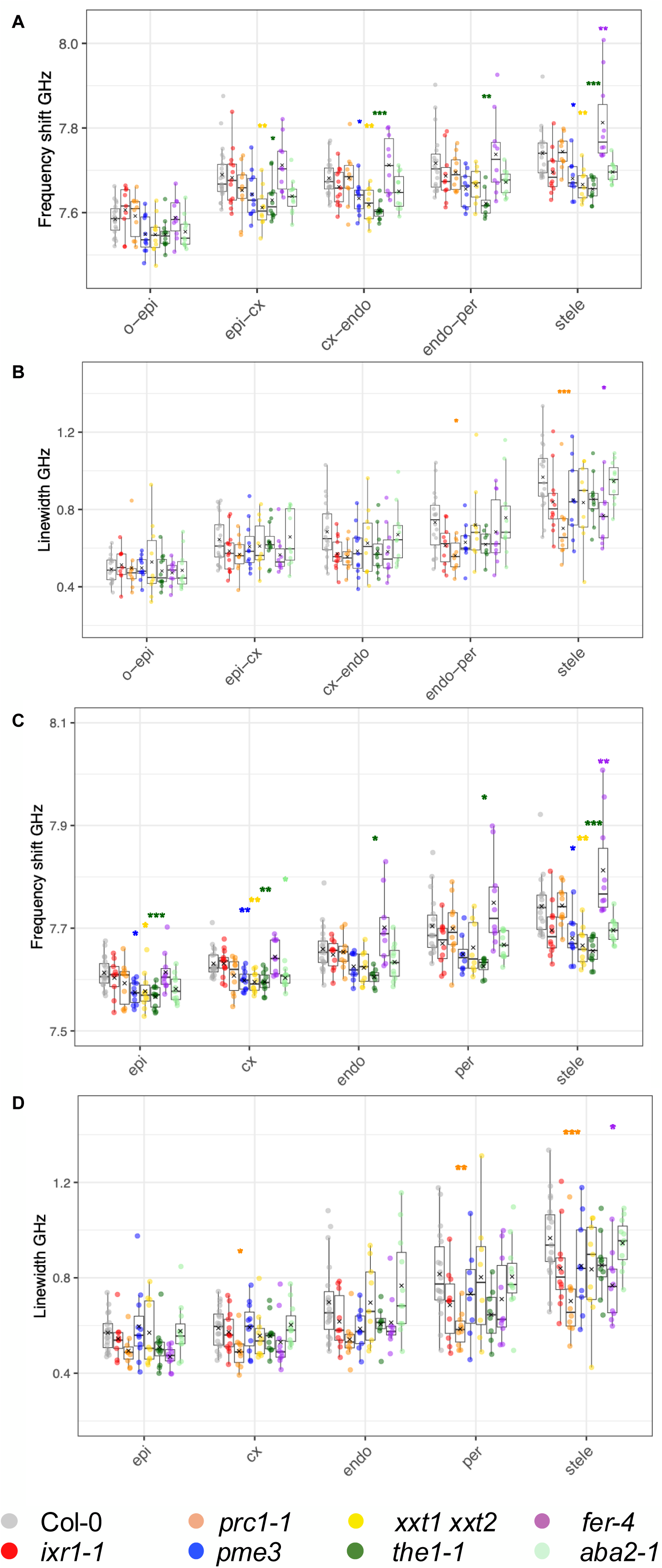
Mechanical properties of mutants affecting cell wall metabolism, cell wall integrity signaling or ABA biosynthesis. Quantification of the mechanical properties in the main root tissues of selected mutants. (A) Frequency shift and (B) linewidth of cell walls. (C) Frequency shift and (D) linewidth of cytoplasm. O = outside, epi = epidermis, cx = cortex, endo = endodermis, per = pericycle.

Surprisingly, these results suggest that impaired cellulose crystallinity or biosynthesis alone apparently do not have a major impact on stiffness. However, cellulose biosynthesis seems to be necessary for maintaining viscosity levels in certain tissues. Interestingly, perturbation of THE1 (and to a lesser extent FER) had more pronounced effects on the mechanical properties than mutations affecting particular cell wall biosynthesis genes. These receptor kinases might participate in multiple signaling pathways in a context-dependent manner, leading to broader mechanical effects when their function is disrupted. The increased frequency shift and decreased linewidth in *fer-4* roots may be explained by a low hydration state of its cell walls, consistent with a mechanical transition from a liquid-like to a solid-like state (*21*). The results for *the1-1*, *pme3* and *xxt1xxt2* suggest that the mechanical properties of the cortex cell layer are particularly sensitive to defects in pectin modification, xyloglucan biosynthesis and CWI maintenance. Intriguingly cytoplasmic and wall stiffness of *the1-1*, *pme3* and *xxt1xxt2* cortex cells is also reduced implying that changes in mechanical properties of cortex cell walls lead to adaptive changes in the cytoplasm, which do not require THE1-mediated signaling. While the expression patterns of the respective genes range from broad (*XXT1*, *PME3*) to more specific (*THE1*), the mutant seedlings exhibited distinct tissue-specific effects on mechanical properties. This implies that tissue layers possess unique mechanical cell wall and/or cytoplasmic properties and that the encoded enzymes have tissue-specific functions, which are not necessarily regulated on the transcriptional level.

### Mechanical properties in the elongation zone change rapidly after stress

Previously, we showed that impairing cell wall integrity with isoxaben (ISX, a cellulose biosynthesis inhibitor) and turgor pressure reduction by sorbitol leads to decreased frequency shifts after 6 hours in Arabidopsis roots compared to controls (*50*). We did not observe differences between the combined treatment (ISX + sorbitol) and controls, suggesting that the individual effects were neutralizing each other. However, these studies lacked the spatial resolution necessary to detect tissue-specific changes in mechanical properties. Here we investigated with higher resolution the mechanical properties of individual tissue layers in roots 3 hours after treatment (Fig. 5 and Supp. Fig. 5), when the effects on cell morphology start to be visible and sorbitol-induced ABA production is at its maximum in our model system (*50*, *51*). The outer epidermal wall did not exhibit frequency shift changes in response to any treatment (Fig. 5A). The epidermis-cortex and stele walls showed decreased frequency shifts in response to sorbitol and ISX + sorbitol treatments, while cortex-endodermis cell walls exhibited reduced frequency shifts in all treatments. In the cytoplasm, sorbitol and ISX + sorbitol treatments significantly reduced the frequency shift in all cell layers, whereas ISX alone only produced a reduction in the cortex. Linewidth values of cell walls and cytoplasm remained unchanged in response to all treatments. Interestingly, the epidermis was least affected by the treatments, suggesting that the outermost tissue perceives and/or responds to stress differently. Furthermore, sorbitol and ISX + sorbitol treatments resulted in similar values for both frequency shift and linewidth within every tissue, suggesting that sorbitol is the primary driver for the changes in mechanical properties. Since frequency shifts were observed both in cell walls and cytoplasm while linewidths did not change, stiffness and viscosity respond apparently differently to CWI impairment and hyper-osmotic stress. Importantly, the findings also show that the mechanical properties of cell walls change *in vivo* in response to osmotic stress in a tissue-specific manner.

**Fig. 5.**
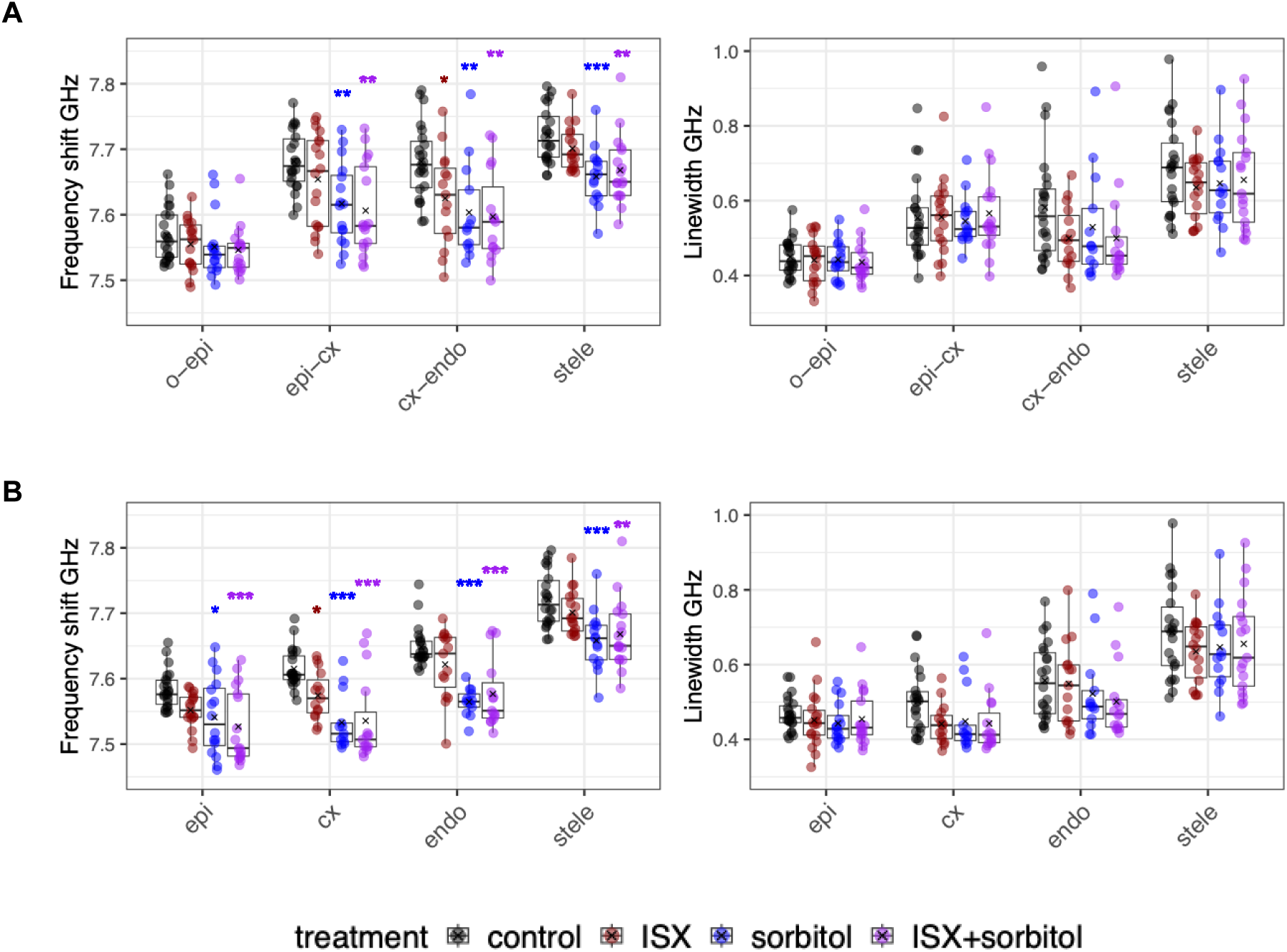
Mechanical properties of cell walls and cytoplasm in the elongation zone in response to different stresses. Mechanical properties of (A) cell walls and (B) cytoplasm. Frequency shift (left) and linewidth (right). ISX = isoxaben. O = outside, epi= epidermis, cx = cortex, endo = endodermis.

## Discussion

We have generated a high-resolution *in vivo* map of mechanical properties of the Arabidopsis seedling root tip using Brillouin microscopy to quantify frequency shift and linewidth (as proxies for stiffness and viscosity) and complemented this with porosity measurements using BODIPY-based molecular rotors. We found that the mechanical properties of different cell layers change along their respective developmental axes, highlighting how properties can change in a short period of time in a highly dynamic manner. Intriguingly, the outer epidermal cell wall exhibited consistently lower frequency shifts than internal cell walls, implying it is not the stiffest cell wall in the root tip. Our results suggest that its mechanical properties may be regulated in a context dependent manner to accommodate specific requirements arising during growth and development (10). Intriguingly, we observed that stiffness changes in cortex cell walls during differentiation to support directional cell elongation in a manner previously postulated (8).

Importantly our *in vivo* studies detected variability in mechanical properties of cell walls surrounding root hairs and individual cortex cells during differentiation. This reinforces the notion that the mechanical properties of cell walls are not always uniform and highlights how we can investigate the coordination between cell wall mechanics and biochemical processes responsible for cell wall homeostasis *in vivo* using Brillouin microscopy. Once, the different cell types are fully differentiated only the stele continues to exhibit pronounced differences. The stele actually displayed already during differentiation the highest stiffness and viscosity, likely reflecting compressive forces exerted from surrounding tissues (*52*). Targeted genetic manipulation of cell wall metabolism (specifically xyloglucan and pectin methylesterification but not celloluse) or signaling induced tissue-specific changes of mechanical properties in the elongation zone exemplified by cortex cells. Intriguingly we observed simultaneously changes in stiffness of cortex cell walls and cytoplasm but not viscosity. Similar effects were observed in seedling roots exposed to hyper-osmotic stress for three hours. This implies that mechanical properties are modified in a coordinated (cell wall-cytoplasm) manner on the subcellular level if cell wall homeostasis is disturbed. These changes may serve as cues for induction of biochemical responses and provide also mechanical feedback to adjacent tissues, but this remains to be tested. Notably, stiffness and viscosity do not always correlate, highlighting that different mechanical states can exist simultaneously, that properties can change in a context dependent manner and that it is important to monitor multiple mechanical properties simultaneously. More importantly, the results illustrate how it is now possible to dissect the mechanical behavior of living cells and tissues *in vivo* in a non-invasive and systematic manner. Integration of information about mechanical properties and biochemical/cell biological processes will be an essential prerequisite to characterize the mechanisms coordinating changes in mechanical properties, biochemistry and cell biology active during plant growth, development and adaptation to a changing environment.

## Acknowledgments

We gratefully thank Benoit Landrein, École Normale Supérieur de Lyon; Chris Sommerville, University of California, Berkeley; Cyril Zipfel, University of Zürich; Timo Engelsdorf, University of Marburg; Herman Höfte, INRAE, Versailles and Malcolm Bennett, University of Nottingham for seeds. We also thank Thomas Dehoux and Jeremie Margueritat, from Université de Lyon 1, and Jitao Zhang, Michigan State University, for help during the setup of the Brillouin microscope.

## Author contributions

Conceptualization: L.A.B., B.T.S., T.H.

Methodology: LA.B., A.B., M.B., J.S.

Investigation: LA.B., F.S., S.M., D.A, M.T.

Visualization: L.A.B., F.S., S.M., D.A., A.I.

Funding acquisition: LA.B., B.T.S., T.H.

Project administration: T.H.

Supervision: L.A.B., B.T.S., T.H.

Writing – original draft: L.A.B., T.H.

Writing – review & editing: All authors

## Data and materials availability

All data are available in the main text or the supplementary materials. Custom MATLAB codes used to calculate the frequency shift and linewidth from the Brillouin images are available upon reasonable request.

## Supplementary Materials

## Materials and Methods

### Plant material

Arabidopsis thaliana seeds of Col-0 accession were obtained from the European Arabidopsis Stock Center in Nottingham (NASC), from laboratories previously publishing them or from collaborators (**Table S1**). Mutant alleles were confirmed via Polymerase chain reaction using the Phire Plant Direct PCR Master Mix Kit (Fisher Scientific F160L). Subsequently, gel electrophoresis or Sanger sequencing (Eurofins Genomics) was performed to confirm the genotype of the plants.

### Plant Growth

Seeds were surface sterilized in three steps: washed in 70% ethanol, then 10 min in 25% (v/v) bleach (Klorin) and then washed them three times with sterile water. Sterile seeds were sown on solid media plates containing 0.5X Murashige & Skoog basal medium (Merck M5519-50L), 0.05% MES sodium salt (Merck M3058-1KG), 1% agar (Merck A1296-1KG), pH 5.75, and stratified 48 hours at 4°C. For Brillouin microscopy experiments the plates were supplemented with 1% sucrose (Merck S7903-5KG). For FLIM experiments, the plates did not containe sucrose. Plants were transferred to a growth chamber and grown under long-day conditions (16 hours light/ 8 hours dark, 150 μmol × m^-2^ × s^-1^ photon flux density, 21°C and 40% humidity) for 6 days except for the early developmental studies where plants were taken from the growth chamber after 4 days.

### Plant Treatments

6-days-old seedlings were transferred to plastic dishes (ThermoFisher Scientific Sterilin 10655821) with control or treatment solutions for three hours. Immediately after, seedlings were mounted on microscopy slides for imaging. Control samples were treated with 0.5X Murashige & Skoog basal medium, 0.05% MES sodium salt, pH 5.75. Treatments were done separately with 600nM isoxaben (Merck 75772-50MG), 300μM sorbitol (D-sorbitol Merck 56021-5KG) or a combined treatment with both compounds.

### Confocal Microscopy

Fluorescent and brightfield images were obtained with an inverted SP8 Leica Confocal Microscope. Samples were illuminated using a 40x water-immersion objective NA 1.1 (Leica). A 488 nm laser was used to excite the plasma membrane reporter p35S:LTI6b-GFP and fluorescence was collected at 500-550 nm using a hybrid detector and a pinhole of 1 airy unit. Brightfield images were collected simultaneously, and the same illumination parameters were used for the non-fluorescent samples.

### Fluorescence Lifetime Imaging Microscopy (FLIM)

Seedlings were stained with 20 μM CarboTag-BDP (*19*) probe for one hour and then transferred to the mounting media. Roots were examined using a Leica SP8 SMD/MP confocal microscope (Leica Microsystems) equipped with a 25x NA 0.95 water immersion objective. White light laser (WLL) 488 nm was used as an excitation line and fluorescence was captured between 500 and 550 nm. The signal was collected using a single-photon avalanche photodiode (APD) with a gain of 200%. The output was connected to a time-correlated single photon counting (TCSPC) module PicoHarp 300 (Picoquant, Germany) and photons were captured over a period of 136 seconds at a repetition rate of 20 MHz. FLIM analysis was performed using SymphoTime 64 (PicoQuant, Germany), employing an n-exponential tailfit fitting model and with the model parameter set to n = 2. The time range (x-axis) in the generated TCSPC graph was adjusted to start from 0.5 nanoseconds after the instrument response function and end 1 nanosecond before the tail. Subsequently, lifetime maps, representing the mean fluorescence lifetime per pixel in nanoseconds, were generated by adjusting the false color scale according to the lifetime histogram. Additionally, the MosaicJ plugin in ImageJ was used to stitch and generate the image of the whole root (*53*, *54*).

### Quantitative Phase Imaging (QPI)

An inverted Axio Observer Z1 microscope (ZEISS) equipped with a quantitative phase imaging/spatial light interference microscope module was used to determine the refractive index of root hair peripheries. A refractive index of 1.33 was set for water as a reference. Temperature in the room was maintained at 20°C for stability during measurements. A 10x objective with NA 0.3 (EC Plan-Neofluar) was used to illuminate the sample and the CELLVISTA (SLIM edition PhiOptics) software was used to control image acquisition with a Hamamatsu Orca Flash v2 sCMOS camera. The refractive index was obtained from 17 root hairs from different roots.

### Brillouin Microscopy

#### Instrumentation

Brillouin microscopy was performed using a custom-built confocal Brillouin microscopy (Supp. Fig. 6A) based on a two-stage virtually imaged phase array (VIPA) configuration, following the design described in (*55*). Excitation was provided using a 532nm laser (Cobolt, Hübner Photonics). The laser was guided through a half-wave plate, a polarized and a beam expander to a polarized beam splitter. Laser light was further directed to quarter-wave plate before entering the back port of an inverted Leica SP8 confocal microscope. Samples were illuminated using a 40x water-immersion objective NA 1.1 (Leica) and the scattered light was collected using a backscattered geometry (180 degrees). Scattered light passed again through the quarter-wave plate and polarized beam splitter before being directed to a 4x NA 0.10 objective (Olympus) which focused the light into a fiber coupler connected to a single mode optical fiber (Thorlabs). The spectrometer consisted of two-stage VIPA etalons (OP-6721-3371-2, 500-600 nm, 30 GHz free spectral range - Light Machinery), vertical and horizontal physical masks to block the elastically scattered light (Raileigh peaks) and a Lyot stop (SF) (*56*)), providing a spectral contrast of 75-80 dB (*55*). An ORCA-Quest qCMOS camera (C15550-20UP, Hamamatsu Photonics) was used to record the Brillouin spectra from the second and third light mode. The microscopy room was maintained at 20°C and the laser was mounted on a heated plate set to 37°C (according to manufacturer recommendations). During Brillouin spectra recording, the confocal microscope stage movement and image acquisition was controlled using the HCImage software (Hamamatsu Photonics). To check for perturbations and phototoxicity, samples were inspected by transmitted-light wide-field illumination before and after the Brillouin spectra was recorded. No visible changes were observed.

#### Imaging

Image acquisition time was 100 ms per pixel and laser power 10-18mW. Image dimensions, frame integration and stage step size were adjusted depending on the image type: 30×200 pixels, 2 frames and 5 μm for root tips; two frames and 1μm for the different developmental regions in the cortex layer and root hairs (variable pixel dimensions); and 100×10 pixels, 4 frames and 1μm for the elongation zone (4- and 6-days-old seedlings, mutants and treatments).

#### Spectral Resolution

According to the manufacturer specifications (Cobolt), the laser linewidth is < 1 MHz. The spectral resolution of the spectrometer, which defines the ability to distinguish light from different wavelengths (*57*), was determined by measuring the width of an elastically scattered peak, from the same orders (between second and third order) as the ones used to measure samples, reflected from a flat mirror (Supp. Fig. 6B). The detected laser peak (blue circles) was fitted with a Lorentzian function (red curve). The linewidth (full width half maximum, FWHM) of the fitted peak was 33.2 pixels, corresponding to 488 MHz, after converting pixels to Hz using water and methanol as calibration samples. We calculated the finesse as the ratio between the free spectral range and the FWHM and obtained a value of 64.

#### Spatial Resolution

The spatial resolution of a Brillouin microscope depends on the attenuation length of the collective excitations (phonons), which is sample dependent (*58*). Scanning across the interface between polymethyl methacrylate and immersion oil type F (Leica, refractive index *n* = 1.518) was performed to quantify the spatial resolution. Data along the interface was fitted with an *erf* function and its derivative was obtained to calculate the FWHM. A lateral resolution of 2 μm and an axial resolution of 8 μm was obtained (Supp. Fig. 6C and D).

#### Spectral Precision

To determine how accurately and reproducibly our frequency measurements are, we repeatedly measured the frequency shift and linewidth of distilled water (Supp. Fig. 6E and F) and calculated the standard deviation from our measurements. The data is represented as a histogram and fitted with a Gaussian curve. We obtained a precision of 10 MHz for the frequency shift and 20 MHz for the linewidth using 10 mW laser power.

#### Image analysis

A custom MATLAB (Mathworks) script was used to extract the Brillouin peaks position and linewidth (FWHM) of the Brillouin peaks (Supp. Fig. 7) after background subtraction and fitting the peaks with a Lorentzian function:

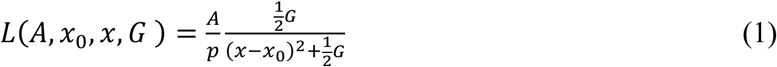

Where A is the intensity, x_0_ is the center of the function, x is a variable representing the position and G is a parameter that specifies the width of the function. We performed deconvolution by subtracting the FWHM from the Raileigh peak (also assumed to have a Lorentzian shape) from the measured FWHM.

Peak positions obtained from distilled water and methanol served as reference samples and were used to calibrate the frequency axis by obtaining the pixel to frequency equivalency and FSR. Calibration was performed using the values of water (*v*_B_ = 7.45 GHz) and methanol (*v*_B_ = 5.45 GHz) with a 532 nm laser wavelength.

Regular measurements of water and methanol were performed and interpolation between measurement time points was used to assign the corresponding calibration values.

Calculated frequency shift and linewidth values were ordered according to image dimensions to generate the corresponding heatmaps. Image overlays between confocal images and heatmaps were done manually in Adobe Illustrator, which were subsequently used for defining the region of interest for quantification.

#### Calculation of the mass density from refractive index

The refractive index data of root hair peripheries obtained using QPI was used to estimate the mass density. The sample was considered as a two-substance mixture of dry and fluid fraction, and the mass density was calculated as (*39*):

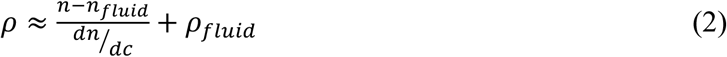

where *ρ* is the mass density of the sample, *n* is the refractive index of the sample, *n_fluid_* is the refractive index of the fluid fraction (refractive index of distilled water *n* = 1.33), *dn/dc* is the refraction increment (∼0.145 ml/g for carbohydrates, (*59*)) and *ρ_fluid_* is the mass density of the fluid fraction (1 g/ml for water).

#### Calculation of the longitudinal modulus

The longitudinal modulus is composed of two terms (*17*, *20*): the real part *M’* (longitudinal storage modulus), which accounts for the elastic behavior, and the imaginary part *M’’* (longitudinal loss modulus), which accounts for the sample’s attenuation properties, reflecting on its viscous behavior.

The frequency shift serves as a proxy for the elastic behavior of a material as it relates to the longitudinal modulus of by the formula (with a scattering collection angle of 180°):

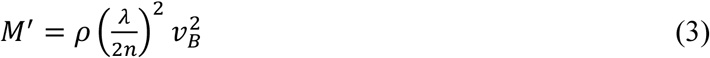

Where *v*_B_ is the frequency shift, *n* is the refractive index, *α* is the excitation laser frequency, *M*’ is the real part of the longitudinal modulus, *ρ* is the material density.

The linewidth is used as a proxy for the longitudinal viscosity of a material. They are related by the formula:

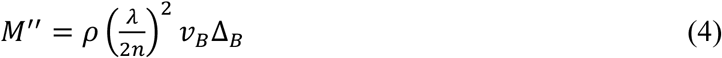

Where *v_B_* is the frequency shift, *n* is the refractive index, *α* is the excitation laser frequency, *M*’’ is the imaginary part of the longitudinal modulus (loss modulus), and *ρ* is the material density. The longitudinal viscosity includes the effects of both shear and bulk viscosity (*17*, *57*).

### Data processing, visualization and statistical analysis

The data was compiled with R and visualized employing the ggplot2 package. Data normality was tested using the Shapiro-Wilk method (α = 0.05).

#### Root tip (Brillouin microscopy)

Data was collected as a list of values starting at the position above the stem cell niche and ending at the elongation zone for every sample. Each value in the list representing the mean of 5 pixels at each position along the root tissue, ensuring that each pixel set contains values for cell walls and cytoplasmic regions in all root zones. Individual data points, representing the mean of all samples (mean of the lists values) at each position, were plotted for all tissues, starting from the position above the stem cell niche (Fig. 1C). Data points were fitted using a generalized additive model (GAM) visualized as solid curves with standard error. A solid vertical line indicates the position of the transition zone based on the first elongated cell in the cortex file.

Between 9 and 12 roots were measured for each tissue.

#### Root tip (FLIM)

After analysis was completed in SymphoTime 64 (PicoQuant, Germany), lifetime data was saved and imported as a text image into ImageJ. Cell walls of individual tissue layers were selected with a segmented line, and a list of the longitudinal lifetime values was obtained using the PlotProfile function. After averaging the values for each tissue, the data was fitted using loess smoothing, visualized as solid curves with standard error. A solid vertical line indicates the position of the transition zone based on the first elongated cell in the cortex file. The cortex, endodermis and stele tissues of 10 roots were analyzed.

#### Longitudinal and transversal cell wall measurements in the cortex (Brillouin microscopy)

For each measured cell, data was collected by selecting a trajectory with the maximum pixel value along the two longitudinal or two transversal cell walls. Data points represent the mean value of each cell wall type (Fig 2C and E). Boxplots were used to visualize the data where measurements from a single cell (longitudinal and transversal cell walls) were paired. A paired t-test was used to evaluate significant differences between the measurements of different cell wall orientations. Between 8 and 11 cells were measured for each developmental zone.

#### Longitudinal and transversal cell wall measurements in the cortex (FLIM)

After analysis of an image was completed in SymphoTime 64 (PicoQuant, Germany), entire longitudinal or transversal cell walls were selected using the free selection tool. The exponential fit and lifetime values were recalculated for the region of interest. Boxplots were used to visualize the data where measurements from a single cell (longitudinal and transversal cell walls) were paired. A paired t-test was used to evaluate significant differences between the measurements of different cell wall orientations. 10 cells were measured for each developmental zone.

#### Elongation zone

Data was collected by selecting a trajectory of pixels with the maximum values along cell walls and calculating the mean (10 pixels for each root, Fig. 3 D and E). Pixels between adjacent cell walls trajectories were used to calculate the cytoplasmic values. Because cell wall and cytoplasm regions in the stele were indistinguishable, the whole stele region was selected to calculate the mean frequency shift and linewidth values.

Box plots were used to represent paired data points of the different root tissues. A repeated measures analysis of variance (ANOVA) test with Holm correction was used to estimate significant differences among tissues. 10 roots were measured for each tissue.

#### Mutants

Data was collected and visualized (Fig. 4) in the same manner as for the elongation zone data. ANOVA with Dunnet’s post hoc test was performed to determine significant differences in mutant vs Col-0 (reference group) measurements for each tissue. Between 9 and 11 roots were analyzed for each mutant line. 19 roots were analyzed for Col-0.

#### Treatments

Data was collected and visualized (Fig. 5) in the same manner as for the elongation zone data. ANOVA with Dunnet’s post hoc test was performed to determine significant differences in treated vs control (reference group) measurements for each tissue. Between 15 and 23 roots were analyzed.

#### Root hairs

Width and length of root hairs are visualized in Supp. Fig. 6B. Each dot represents an individual root hair from a different root. To quantify the mechanical properties, root hair cells were divided into base, middle and tip regions. High-value pixels of ellipsoidal regions at the base, likely belonging to nuclei, were excluded from the analysis. Each root hair was represented as a set of three paired data points, one in each region (Supp. Fig. 2E). A repeated measures ANOVA test with pair-wise t-test and Bonferroni correction was used to determine significant differences among the root hair regions. Quantification was done in 19 root hairs from different roots.

Statistical on p-values are reported as ns > 0.05, * < 0.05, ** < 0.01, *** <0.001.

---

**Table S1.** Arabidopsis lines used in this study.

| AGI code | Genotype | Background | Source |
| --- | --- | --- | --- |
| - | Col-0 |  | NASC |
| - | p35S-LTI6b-GFP | Col-0 | Benoit Landrein<br>(École Normale Supérieure Lyon) |
| At5g05170 | *ixr1-1* | Col-0 | Chris Sommerville<br>(UC Berkeley) |
| At5g64740 | *prc1-1* | Col-0 | Cyril Zipfel<br>(University of Zürich) |
| At3g14310 | *pme3*<br>N400106 | Col-0 | Timo Engelsdorf<br>(University of Marburg) |
| At3g62720<br>At4g02500 | *xxt1 xxt2* | Col-0 | NASC |
| At5g54380 | *the1-1* | Col-0 | Herman Höfte<br>(INRAE Versailles) |
| At3g51550 | *fer-4* | Col-0 | Malcolm Bennett<br>(University of Nottingham) |
| At1g52340 | *aba2-1* | Col-0 | (60) |

**Supp. Fig. 1.**
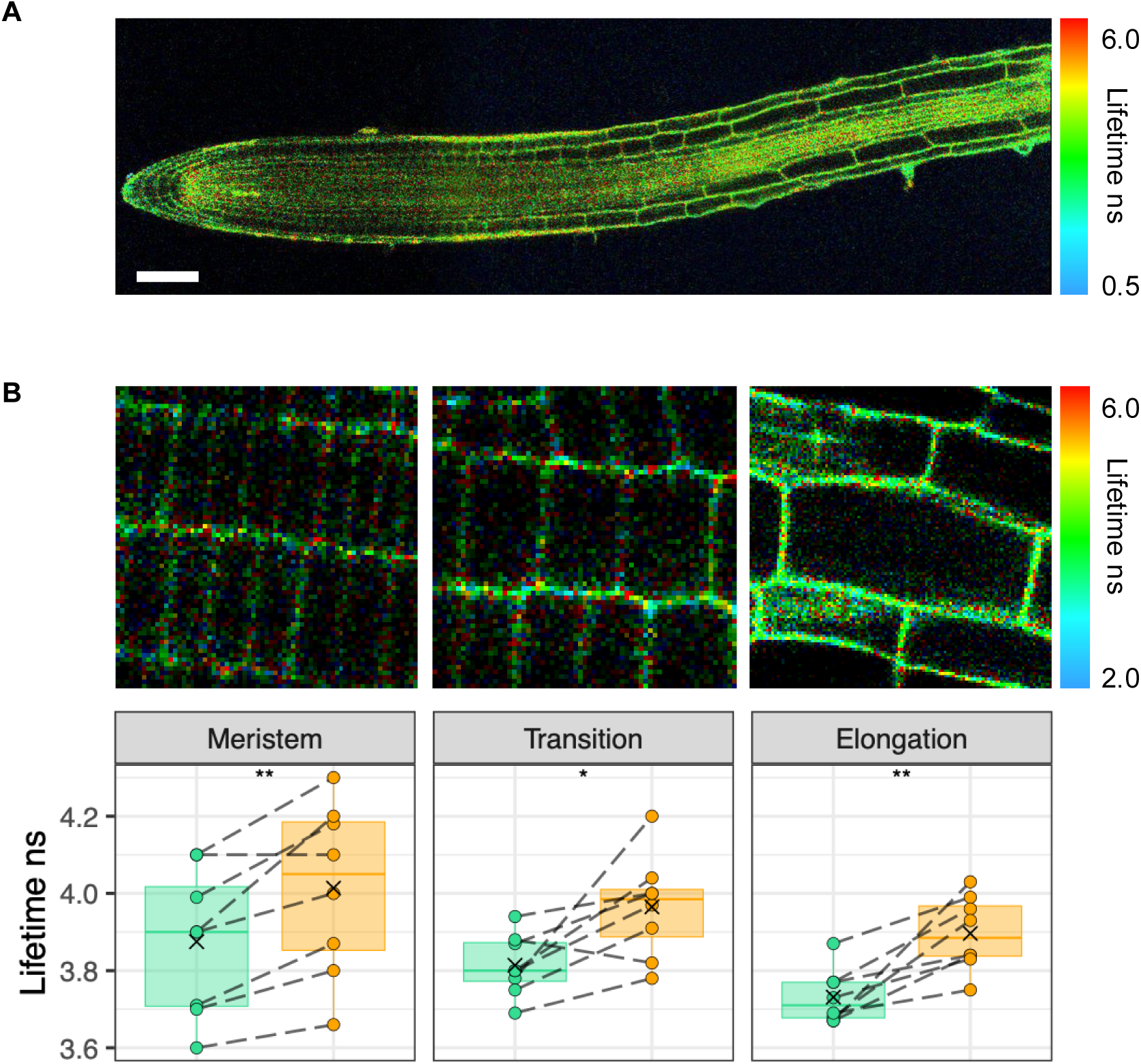
Measurement of cell wall porosity using CarboTag-BDP molecular rotors. Fluorescence lifetime images of (A) the root tip (scale bar 100 *μ*m) and (B) distinct developmental zones along the cortex layer. (Bottom) Quantification of the fluorescence lifetime in longitudinal (green data) and transversal cell walls (orange data) in every zone.

**Supp. Fig. 2.**
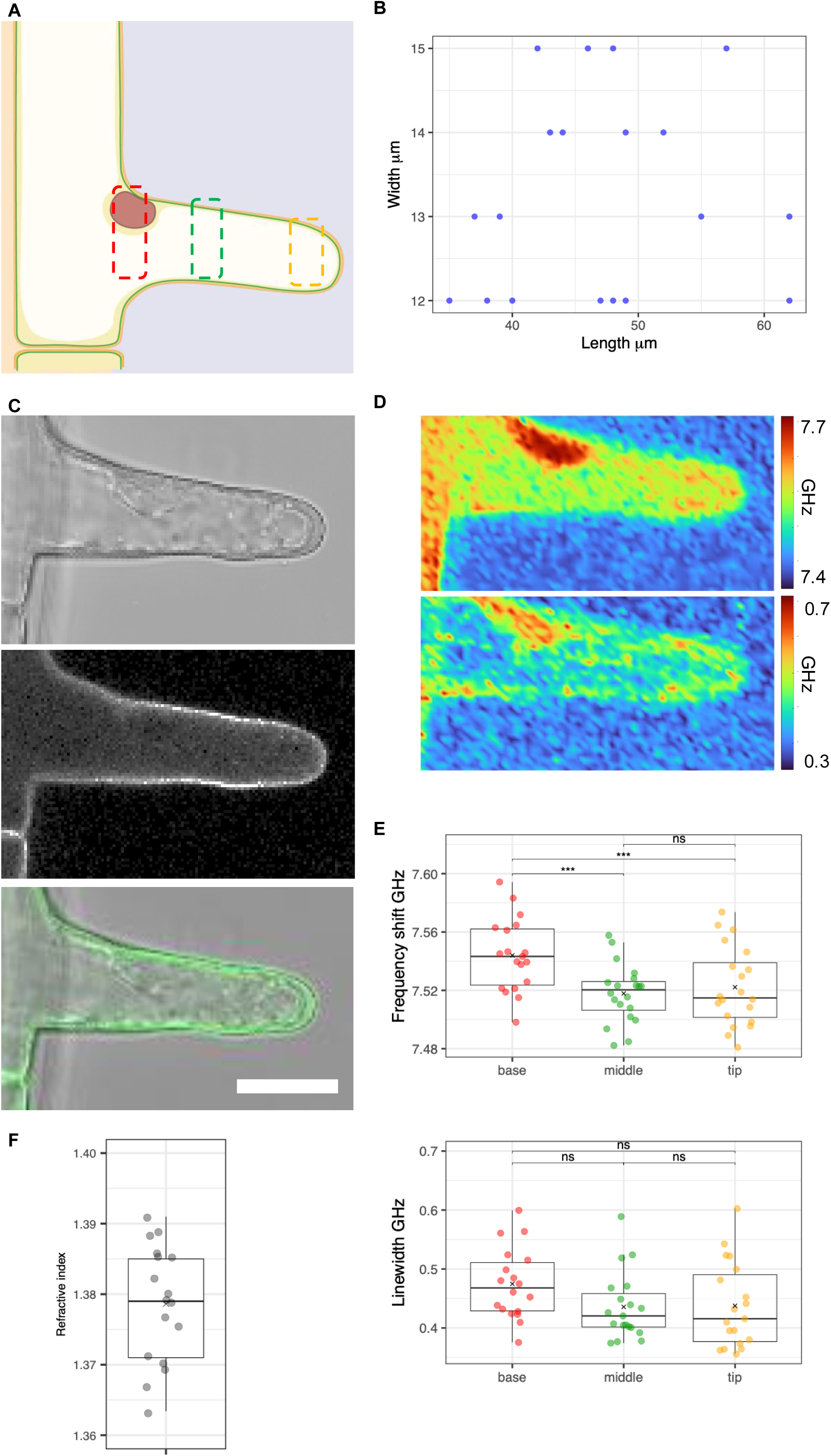
Mechanical properties of root hairs. (A) Schematic representation of a developing root hair. Regions used for quantification are marked by dashed rectangles. (B) Length and width of analysed root hairs Every blue dot represents a root hair from a different root. (C) Brightfield (top), confocal image with p35S:LTI6b-GFP plasma membrane marker (middle), overlap (bottom). Scale bar 20 µm. (D) Heatmaps (35 x 70 µm) of the frequency shift (top) and linewidth (bottom). (E) Quantification of frequency shift (top) and linewidth (bottom) at base, middle and tip regions of root hairs. (F) Quantification of the refractive index at root hair peripheries.

**Suppl. Fig. 3.**
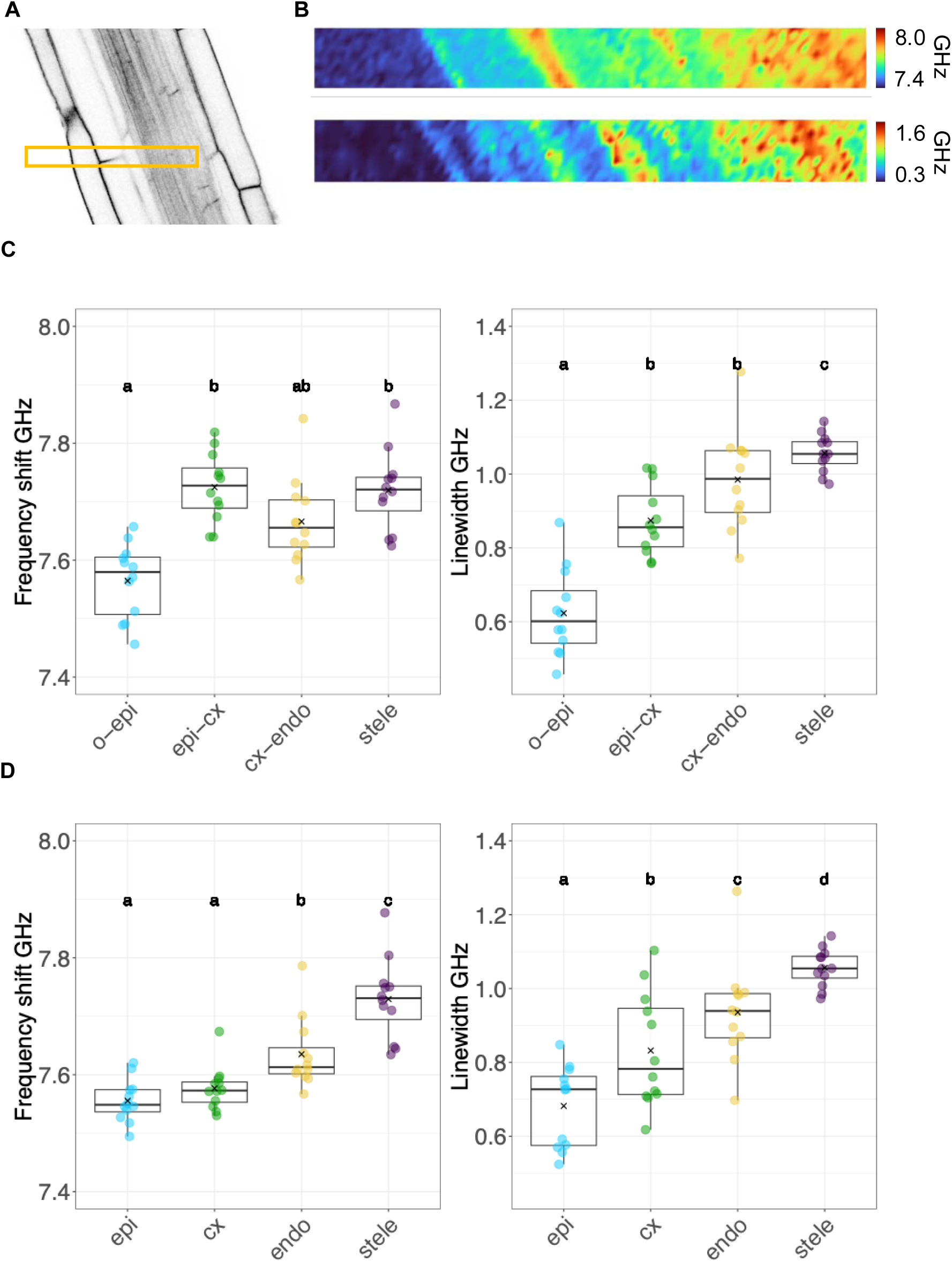
Mechanical properties in the elongation zone of 4-days-old seedlings. (A) Confocal image of the root elongation zone with p35S:LTI6b-GFP. (B) Heatmaps of the mechanical properties in the elongation zone tissues (10 x 100 µm), frequency shift (top) and linewidth (bottom). (C) Quantification of the mechanical properties in cell walls. (D) Quantification of the mechanical properties in cytoplasm. (C-D) Frequency shift (left) and linewidth (right). O = outside, epi= epidermis, cx = cortex, endo = endodermis.

**Suppl. Fig. 4.**
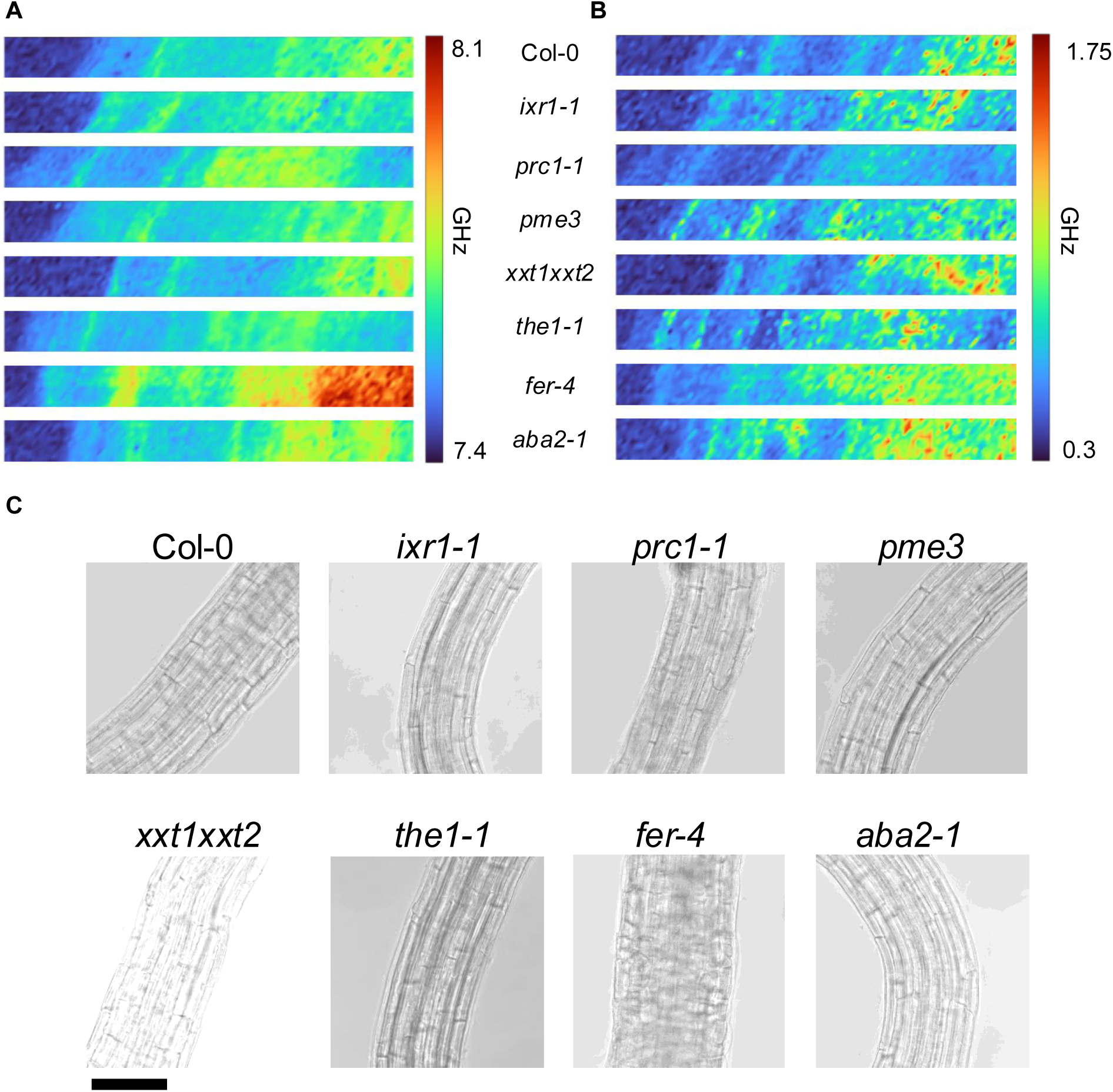
Mechanical properties in the root elongation zone in mutants affected in cell wall metabolism, cell wall integrity signaling or ABA biosynthesis. Heatmaps of (A) frequency shift and (B) linewidth in the selected mutants (10×100 µm). (C) Brightfield images of the elongation zone. Scale bar 100 *μ*m.

**Suppl. Fig. 5.**
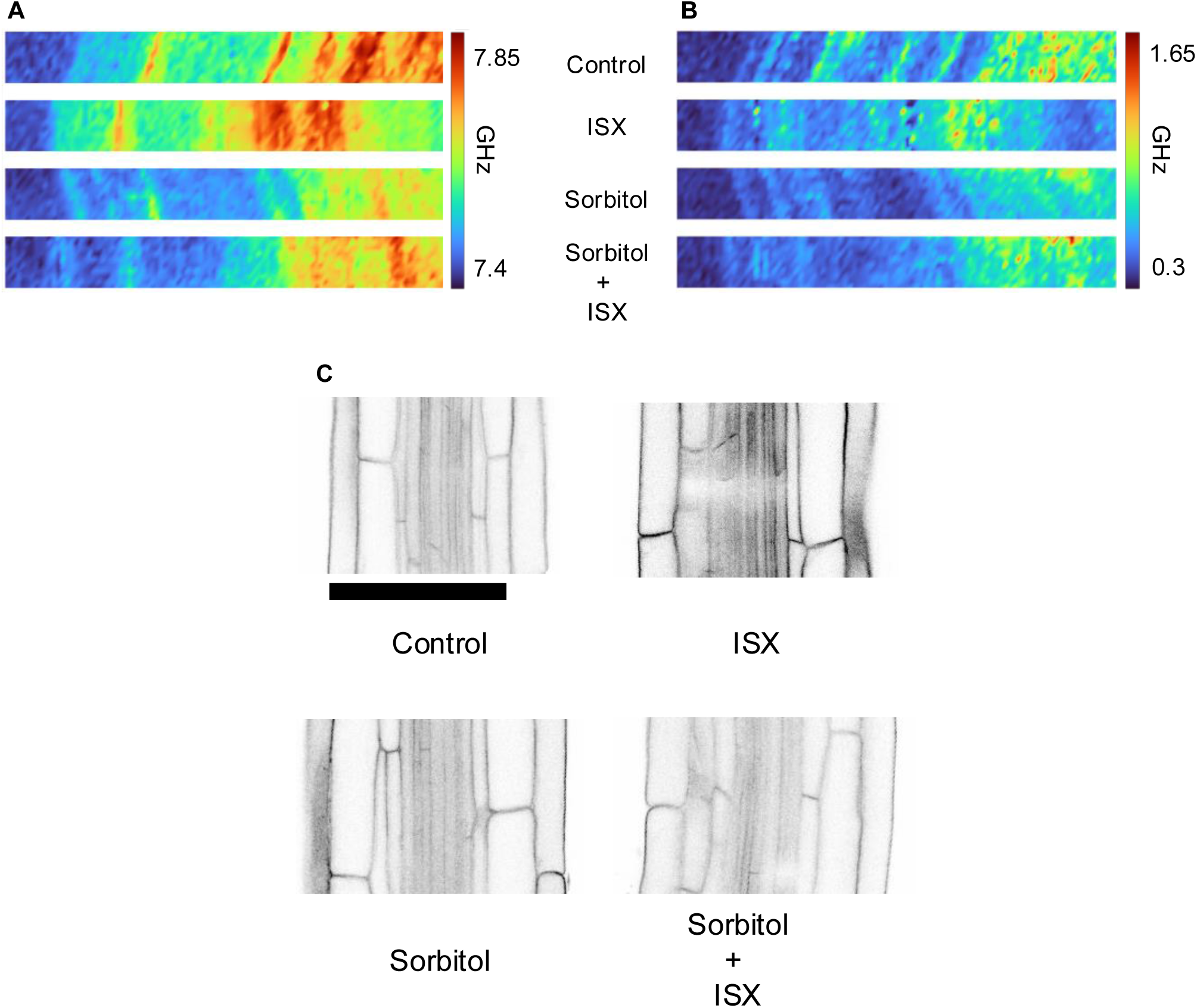
Mechanical properties of tissue layers in the root elongation zone in response to different stresses. Heatmaps (10 x 100 *μ*m) of the (A) frequency shift and (B) linewidth. (C) Confocal images with plasma membrane marker p35S:LTI6b-GFP. Scale bar 100 *μ*m.

**Suppl. Fig. 6.**
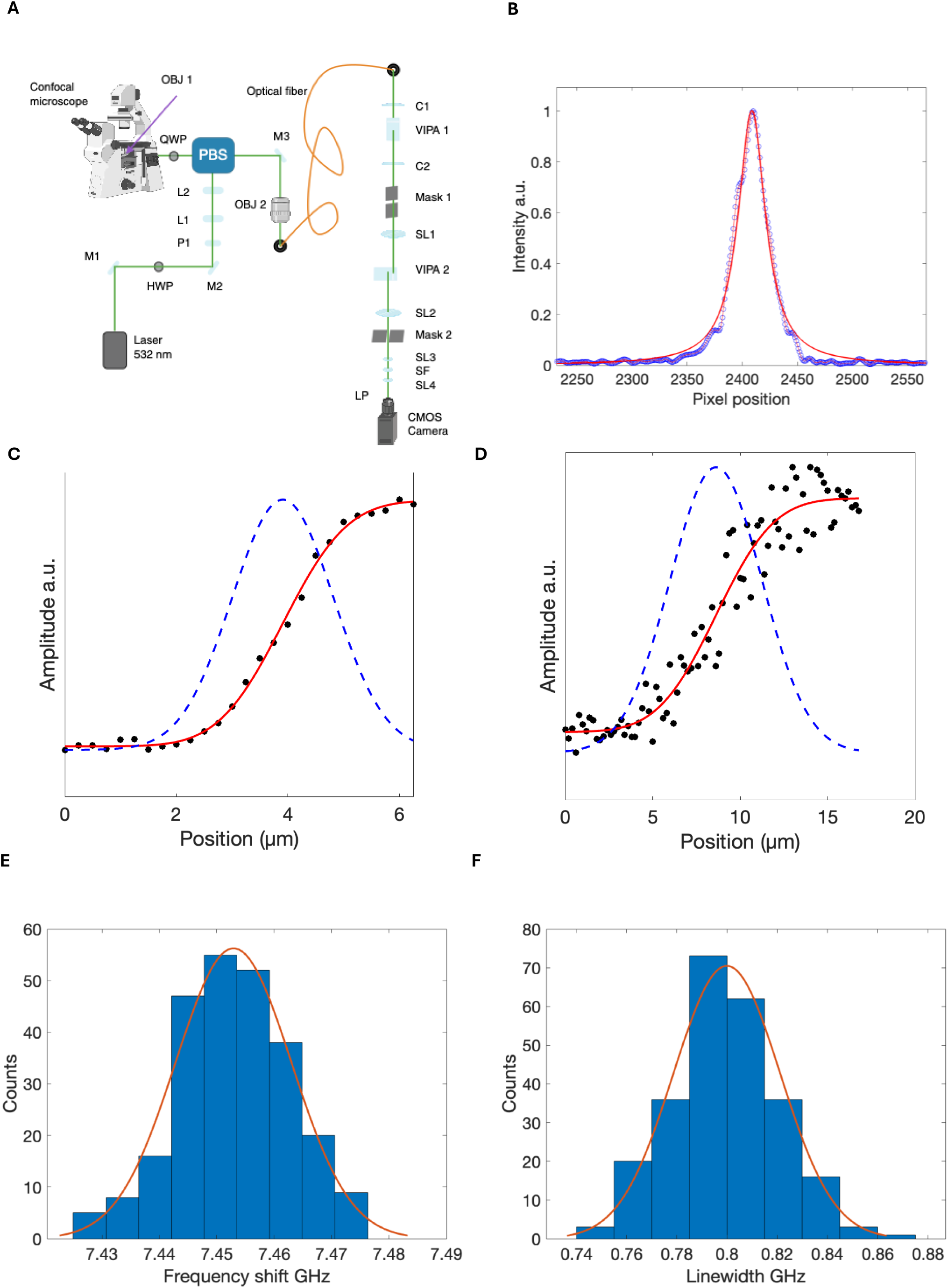
Brillouin microscope instrumentation and resolution. (A) Schematic of the Brillouin microscope setup. M1 = mirror 1, HWP = half-wave plate, M2 = mirror 2, P1 = polarizer 1, L1 = beam expander lens 1, L2 = beam expander lens 2, PBS = polarized beam splitter, QWP = quarter-wave plate, OBJ1 = objective 1, M3 = mirror 3, OBJ2 = objective 2, C1 = cylindrical lens 1, VIPA1 = virtually imaged phased array 1, C2 = cylindrical lens 2, SL1 = spherical lens 1, VIPA2 = virtually imaged phased array 2, SL2 = spherical lens 2, SL3 = spherical lens 3, SF= spatial filter, SP4 = spherical lense 4, LP = lens pair. Created with Biorender. (B) Spectral resolution determination by fitting a Lorentzian function (red curve) to Rayleigh peak data (blue open circles).(C) Lateral and (D) axial resolution measurement across an interface between oil and polumethyl methacrylate. Black dots represent data points, *erf* fit in red and its derivative in dashed blue line. Histogram of repeated (E) frequency shift and (F) linewidth measurements on distilled water to calculate the instrument spectral precision. A Gaussian fit (red curve) was used to calculate the FWHM from the data (blue bars).

**Suppl. Fig. 7.**
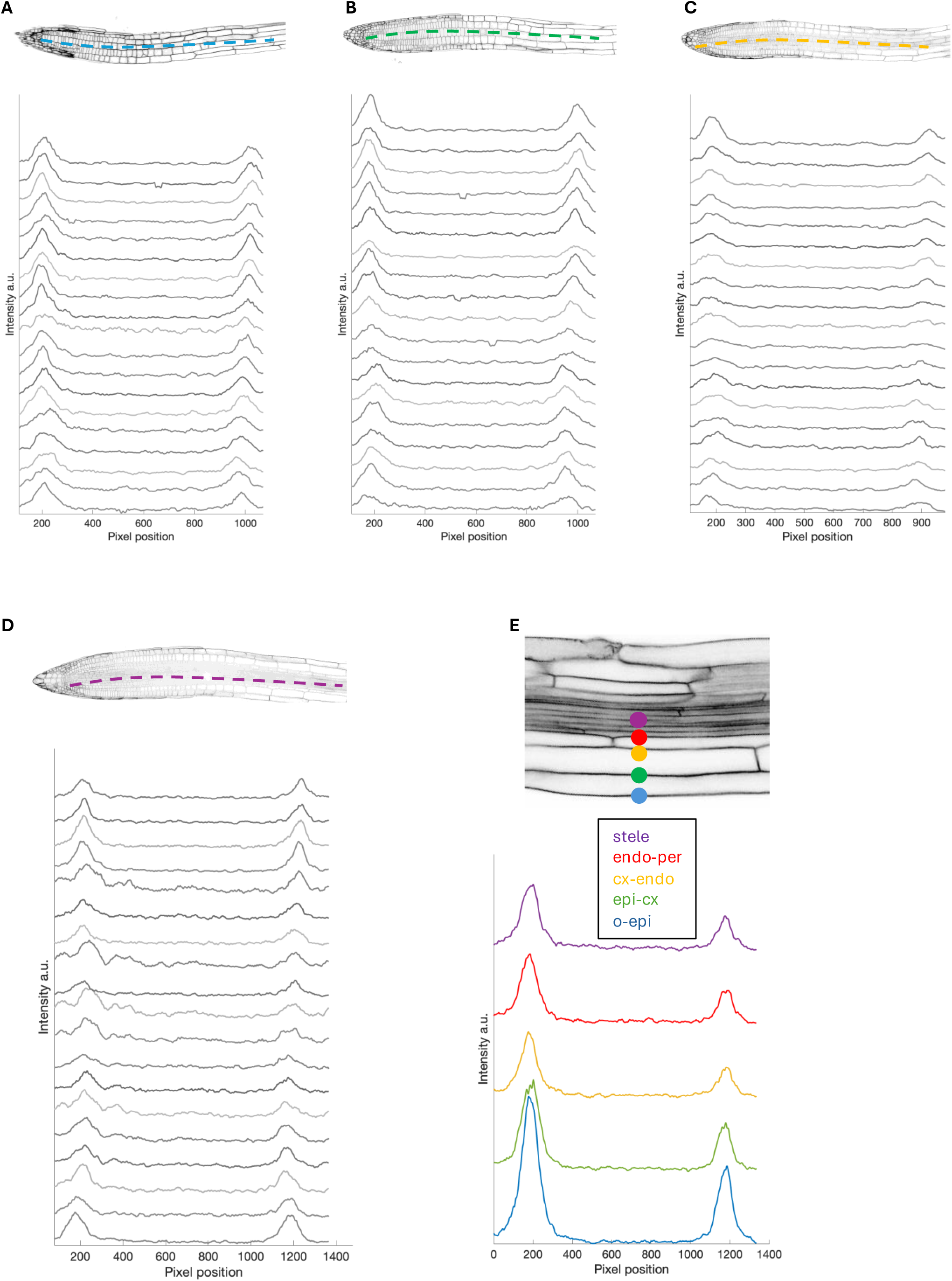
Brillouin spectra from different tissues. The Brillouin spectra were visually inspected to verify the quality of the Brillouin peaks and check for noise levels. For each tissue plane of Fig. 1, spectra located every 20 *μ*m along developmental axes (colored dashed lines in the root) were plotted as representative examples of (A) epidermis, (B) cortex, (C) endodermis and (D) stele tissues. (E) Spectra localized at cell wall positions of the elongation zone (colored dots), corresponding to Fig. 3.

